# Characterization of the vaginal DNA virome in health and dysbiosis: an opening study in patients with non-female factor infertility

**DOI:** 10.1101/755710

**Authors:** Rasmus R. Jakobsen, Thor Haahr, Peter Humaidan, Jørgen Skov Jensen, Witold Kot, Josue Castro-Mejia, Ling Deng, Thomas D. Leser, Dennis S. Nielsen

## Abstract

**Background:** Bacterial vaginosis (BV) is characterised by a reduction in *Lactobacillus* spp. abundance and increased abundance of facultative anaerobes, like *Gardnerella vaginalis*. BV aetiology is not fully understood, but bacteriophages could play a pivotal role causing perturbation of the vaginal bacterial community. Here we investigate the vaginal viral community, including bacteriophages, and its association to the bacterial community and BV-status.

**Methods:** Vaginal samples from 48 patients undergoing IVF treatment for non-female factor infertility were subjected to metagenomic sequencing of purified virus-like particles. The vaginal viral community was characterized and correlated with BV-status, bacterial community structure and presence of key vaginal bacterial species.

**Results:** The majority of identified vaginal viruses belonged to the class of double-stranded DNA bacteriophages, with eukaryotic viruses constituting 4% of total reads. Clear links between viral community composition and BV (q = 0.006, R = 0.26) as well as presence of *L. crispatus* (q = 0.001, R = 0.43)*, L. iners, Gardnerella vaginalis* and *Atopobium vaginae* were found (q < 0.002, R > 0.15). Interestingly, also the eukaryotic viral community was correlated with BV-status (q = 0.018, R = 0.20).

**Conclusions:** The vaginal virome is clearly linked with bacterial community structure and BV-status.

**Clinical Trials Registration:** NCT02042352.

## Introduction

The vaginal microbiota (VMB) refers to the microorganisms inhabiting the vagina. Until now most studies have focused on bacteria and fungi, whereas little is known about the viral community. When dominated by *Lactobacillus (Lb.)* spp., especially *Lb. crispatus*, the VMB has a protective role preventing bacterial vaginosis, yeast infections, sexually transmitted infections (STIs) and urinary tract infections[1]. Bacterial vaginosis (BV) is the most common dysbiosis of the VMB, affecting 10%-30% of reproductive age women in developed nations[2]. BV is characterized by a reduction of vaginal *Lactobacillus* abundance and an increase in the number of other facultative anaerobic bacteria[3]. A dysbiotic VMB and BV have been reported to be an important risk factor for STI acquisition and adverse reproductive outcomes[4]. Most healthy women have a stable and relatively simple vaginal bacterial community dominated by one single *Lactobacillus* spp. However, it is well-known that perturbations of the VMB occurs during menses and antibiotic treatment[5], and is negatively affected by increasing number of sexual partners[6] while being protected by male circumcision[7]. Although *Gardnerella vaginalis* is generally accepted as one of the key bacteria involved in BV pathogenesis[8], the exact etiology of BV is still undergoing further investigation.

The total collection of vaginal viruses (the vaginal “virome”), has only been sparsely investigated, but there is increasing evidence that bacteriophages (or “phages”) are a factor in certain diseases related to gut microbiome dysbiosis[9]. Further, phages adhere to and are significantly enriched in mucosal surfaces, possibly providing what has been termed “non-host derived immunity” against infection[10,11].

Phages are viruses targeting bacteria in a host-specific manner. Phages use one of two fundamentally different ways of replication. Lytic phages infects the bacterial cell, directs the biosynthesis to new phages and then lyses the cell to release new phages. Temperate phages are also able to enter the lytic cycle, but additionally - and in contrast to lytic phages - they are able to replicate through the lysogenic cycle. Here the phage inserts its genetic material into the prokaryote genome as a prophage, which lies latent and is transmitted vertically during bacterial replication until activated whereupon the lytic cycle commences[12].

A possible mechanism underlying VMB dysbiosis could be prophage induction causing community shifts[13]. In support of this hypothesis, it has been shown that VMB-related *Lactobacillus* spp. contain prophages that are activated by environmental stressors such as smoking[14].

Investigation of the virome by metagenomic sequencing can be performed with or without purification of virus-like particles (VLPs). Purification of VLPs has the advantage of removing the majority of bacterial and host DNA allowing deeper sequencing of viral DNA, with the disadvantage that some viral particles may be lost during purification[15]. Alternatively, viral reads can be identified and recovered from metagenomes by matching to known viral genomes, or using a variety of computational tools[16]. This has the disadvantage that the majority of reads from full metagenome sequencing will be from bacterial genomes due to their much larger genome size[17], and significant bias towards detection of viral families more characterized in genome databases[18].

A recent study investigated possible links between vaginal dysbiosis and the risk of HIV acquisition in South African women. Moreover they also investigated the vaginal virome, but reported no distinct viral community structures within their cohort [19]. However, these findings should be interpreted with caution, as the experimental protocol was based on filtration of VLPs from low volumes of cervicovaginal lavage (CVL) without any up-concentration steps, yielding low input for downstream analysis. Furthermore, the bioinformatics analysis was based on reads mapping exclusively to NCBI viral sequences, therefore excluding uncharacterized viruses or phages incorporated into known bacterial genomes as prophages. Another recent study used bioinformatically extracted viral reads from vaginal metagenomes[20], noting the presence of eukaryotic virus families Herpesviridae and Partitiviridae, but without analysing the bacteriophage community in detail. Finally, a study, based on targeted purification of vaginal eukaryotic viral nucleic acids using biotinylated probes targeting all vertebrate viruses prior to high throughput sequencing, found that increased vaginal eukaryotic viral richness is significantly associated with preterm birth [21].

Here we report a cross-sectional investigation of the vaginal virome of 48 Danish women submitted to IVF treatment. Using metagenome sequencing of highly purified VLPs from vaginal swabs, we aimed to elucidate both the overall composition and variability of the vaginal bacteriophage and eukaryotic viral component and explore associations with the vaginal bacterial community state types considered either healthy or dysbiotic.

## Materials and methods

### Patients and samples

This study is a sub-study of a larger study which developed a novel molecular based diagnosis of abnormal vaginal microbiota reporting poor reproductive outcome of IVF treatment[22,23]. The study was approved by the regional ethical committee of Central Denmark Region (file number 1-10-72-325-13) and the Danish Data Protection Agency (file number 1-16-02-26-14). Furthermore, the project was pre-registered at clinicaltrials.gov (file number NCT02042352). All patients signed written informed consent prior to enrollment. The present study involves 48 women admitted to IVF treatment due to either male factor, being single or lesbian. Patients with known female reproductive disorders were excluded. This was done to focus the study on establishing a baseline of the vaginal virome of healthy, reproductive-age women. In brief, patients were recruited at two IVF centers in Denmark and they were prospectively included at their first consultation before initiation of their IVF treatment. Samples were taken from the posterior fornix during speculum examination prior to treatment. One sample was used for Gram staining and Nugent scoring [24]. The second sample was taken for molecular analyses using a 1 ml Copan ESwab™ (Copan Italia, Brescia, Italy) and frozen at −80°C until further use.

### Characterisation of the bacterial component of the VMB

The bacterial component of the VMB was characterised using a combination of 16S rRNA gene amplicon profiling and quantitative (q) PCR analyses for *G. vaginalis*, *A. vaginae*, *L. crispatus*, *L. jensenii*, *L. gasseri* and *L. iners*[23]. These data have previously been published[22] and were made available for the present study.

### Virus like particles (VLP) purification, viral DNA extraction and sequencing

Vaginal material was diluted in 5 ml of SM-buffer (200 mM NaCl, 50 mM Tris • HCl, 8 mM MgSO4 ⋅7H2O, pH 7.4) and filtered through 0.45 µm Minisart^®^ High Flow PES syringe filter (Cat. No. 16533, Sartorius, Germany). The filtrate was concentrated, and particles below 50kDa were removed using Centriprep^®^ Ultracel^®^ YM-50kDa filter units (Cat. No. 4310, Millipore, USA) centrifuged at 1500 x g at 25°C until approximately 300 µL was left in the outer tube. This was defined as the concentrated virome. The 50 kDa filter from the inner tube was removed and added to the concentrated virome and stored for at 4°C until nucleic acid extraction. 140 µL of virome was treated with 2.5 units of Pierce™ Universal Nuclease (Cat. No. 88700, ThermoFisher Scientific, USA) for 3 min, to remove free DNA/RNA molecules. The nucleases were inactivated by 560 µL AVL buffer from the QIAamp^®^ Viral RNA Mini kit (Cat. No. 52904, Qiagen, Germany). For viral DNA extraction the NetoVIR procotol[25] was followed from step 11-27, (with the AVE elution buffer volume adjusted to 30 µL). Samples were stored at −80°C until viral genome amplification. The Illustra Ready-To-Go GenomiPhi V3 DNA Amplification Kit (Cat. No. 25-6601-96, GE Healthcare Life Sciences, UK) was used for viral genome amplification following the instructions of the manufacturer, but with DNA amplification decreased to 60 min to decrease amplification bias[26]. Genomic DNA Clean & Concentrator™-10 units (Cat. No D4011, Zymo Research, USA) were used to remove DNA molecules below 2 kb (following the instructions of the manufacturer). Viral DNA libraries were generated using the Nextera XT DNA Library Preparation Kit (Cat. No. FC-131-1096, Illumina, USA) following the instructions of the manufacturer. Tagged libraries were sequenced as part of a flow cell of 2 × 150 bp pair-ended NextSeq 550 (Illumina, CA) sequencing run.

### Viral-Operational Taxonomic Unit (vOTU) table

Following assembly and quality control (**Supplementary materials S6**), high-quality dereplicated reads from all samples were merged and recruited against all the assembled contigs at 95% similarity using Subread[27] and a contingency-table of reads per Kbp of contig sequence per million reads sample (RPKM) was generated forming the vOTU (viral-operational taxonomic unit)-table. vOTU taxonomy was determined by querying the viral contigs against a database containing taxon signature genes for virus orthologous group hosted at www.vogdb.org using USEARCH-ublast (e-value 10^−3^). The set of vOTUs assigned eukaryotic viral taxonomy was considered the eukaryotic viral component. Sequences are available at European Nucleotide Archive (Data has been submitted to ENA – link will be provided for as soon as available).

### Community analysis

Prior to bioinformatics analysis, vOTU’s which did not have a relative abundance above 0.5% in at least 2 samples were discarded. The sum abundance of removed vOTU’s constituted 3% or less. vOTUs shorter than 3kb were removed. Cumulative sum scaling [28] normalisation was carried out using Quantitative Insight Into Microbial Ecology 1.9.1[29] (QIIME 1.9.1). QIIME 2 (2018.4 build 1525276946)[29] was used for subsequent analysis steps of alpha- and beta diversity statistics. Shannon, Simpson and Richness alpha-diversity and Bray-Curtis dissimilarity index, Jaccard index and Sørensen-Dice coefficient beta diversity matrices were calculated. Wilcoxon Rank Sum Test evaluated pairwise taxonomic differences and Analysis of similarities (ANOSIM) and Kruskal-Wallis tests was performed for group comparisons. For presense/absence of key bacterial species the qPCR the threshold measured by Haahr et al. was used[23]. The k-mer based host-phage prediction algorithm WiSH(1.0)[30] was used to predict prokaryotic hosts based on vOTU sequences. The WiSH host prediction models were based on 9,747 bacterial genomes and 1,970 phage genomes derived from NCBI RefSeq[31], September 2018.

Integrase content was estimated by mapping all viral reads/sample against a database of 247.000 viral integrase and transposase genes UniProt[32], September 2018 followed by calculation of the fraction of mapping reads. This was done to avoid bias caused by incomplete viral genome assembly. Each viral genome is unlikely to contain more than one integrase gene copy[33]. A temperate phage genome with fractured assembly would therefore appear as several vOTUs, with only one containing an integrase and be counted as one temperate and several lytic phages, which is circumvented using the approach outlined above.

Associations between the bacterial and viral community were performed using the mixOmics[34] R package using CSS-normalized OTU-tables as input. Regularized Canonical Correlation Analysis (rCCA) and sparse Partial Least Squares (sPLS) were used to perform combined integration and variable selection on the viral and bacterial OTU-tables. The component tuning cross-validation procedures were double-checked using the shrinkage method for rCCA and leave-one-out cross validation for sPLS.

## Results

### Samples and sequencing

Purified viral particles from vaginal swabs (n=48) were whole-genome sequenced generating a total of 2,674,574 reads (median of 44,868 paired end reads per sample) after joining, trimming, quality control and discarding human and bacterial reads. A total of 773 viral vOTUs were retained after de novo assembly and filtering, sized from 3 to 85 kb with a median of 7.5 kb in length. Basic patient characteristics are shown in Table 1.

**Table 1.** Overview of patient characteristics

| **Table 1.** |  |
| --- | --- |
|  | **Patients, No. (%)<sup>a</sup> (n = 48)** |
| Age, median (range), y | 29 (23-41) |
| Body mass index, median(range)<sup>b</sup> | 31 (17.5-41) |
| Ethnicity |  |
| Caucasian | 83 (40) |
| Eastern European | 8 (4) |
| Other | 6 (3) |
| Asian | 2 (1) |
| Cause of infertility |  |
| Male factor | 58 (28) |
| Single | 13 (6) |
| Lesbian | 3 (2) |
| pH | 4 (4-7) |
| Nugent Group |  |
| BV | 25 (12) |
| Normal | 73 (35) |
| Intermediate | 2 (1) |

<sup>a</sup> Data represent No. (%) of patients unless otherwise specified.
<sup>b</sup> Body mass index is calculated as the weight in kilograms divided by height in meters squared.
<sup>c</sup> Based on Nugent scoring of Gram-stained smears

### Composition of the vaginal virome

A total of 61% of the *de novo* constructed contigs were identified as viral by matching to viral sequence databases. The remaining 39% of vOTUs had no matches in viral databases, but neither did they match bacterial sequences nor human DNA which could be due to the identification of previously unidentified viruses. Viral matches were predominantly from the class of double stranded DNA (dsDNA) viruses, unidentified viruses and a small proportion of single stranded DNA (ssDNA) viruses (Figure 1, **Figure S1**). The investigated samples have previously been classified as either BV-positive, BV-negative or intermediate based on Nugent scoring of the bacteria in a Gram stained smear [23].There was no significant difference in viral alpha diversity between BV-positive and BV-negative samples as determined by neither Shannon diversity index nor number of observed vOTUs (**Figure S1B, C**). Similarly, no correlation was found between viral and bacterial alpha diversity (**Figure S1D**). Eukaryotic viruses were identified in all samples constituting on average 4% of total reads (**Figure S2A**).

**Figure 1.**
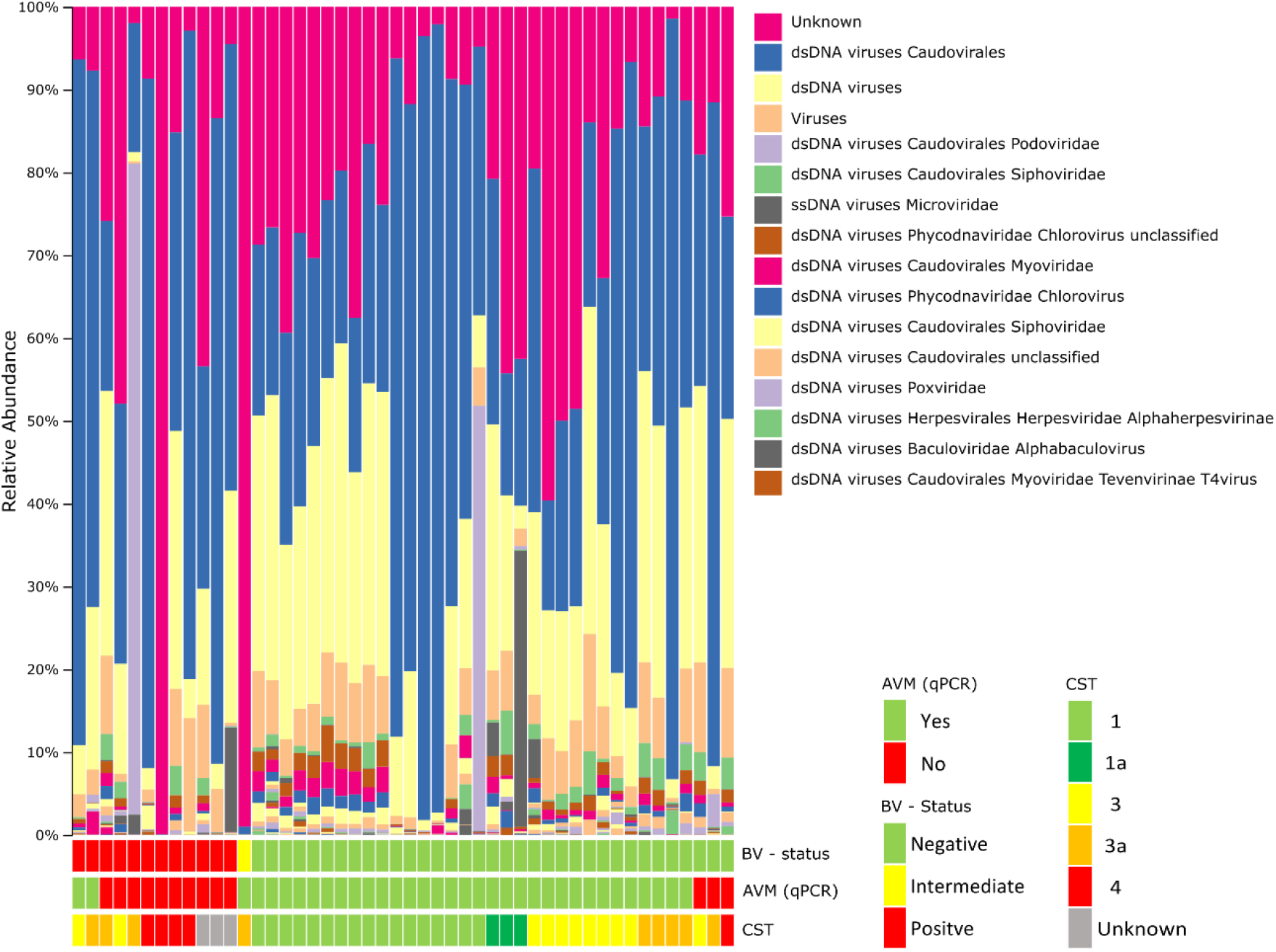
Viral community composition by relative abundance, grouped by bacterial vaginosiss (BV), abnormal vaginal microbiota (AVM) and community state (CST) of sample bacterial community. Taxonomy based on viral database sequence matching.

### Viral beta diversity is significantly correlated with BV-status

The viral community composition (as determined by Bray-Curtis dissimilarity metrics) was significantly different between BV-positive and BV-negative women (q = 0.006, R = 0.26) (Figure 2A). Comparison with a re-analysis of previously published data[22] on the bacterial community of the same samples (based on 16S rRNA gene amplicon data) showed that the vaginal virome composition is as strong a predictor of BV (Nugent score-based diagnosis) as the bacterial composition (q = 0.003, R = 0.50) (F**igure S3**). qPCR-based quantification of key bacteria has previously been shown to be a more accurate predictor of BV than traditional Nugent scoring[35]. In accordance, we found that virome composition corresponded strongly with the qPCR determined presence/absence of *L. crispatus* (Figure 2B) and *L. iners* as well as the BV-associated *Gardnerella vaginalis* and *Atopobiom vaginae* species (**Table S2D**). The presence of *L. gasseri* and *L. jensenii* were on the other hand not significantly associated with the viral component. Interestingly, the composition of the eukaryotic viral community also varied with BV-status (q = 0.018, R = 0.20) (**Figure S2B**).

**Figure 2.**
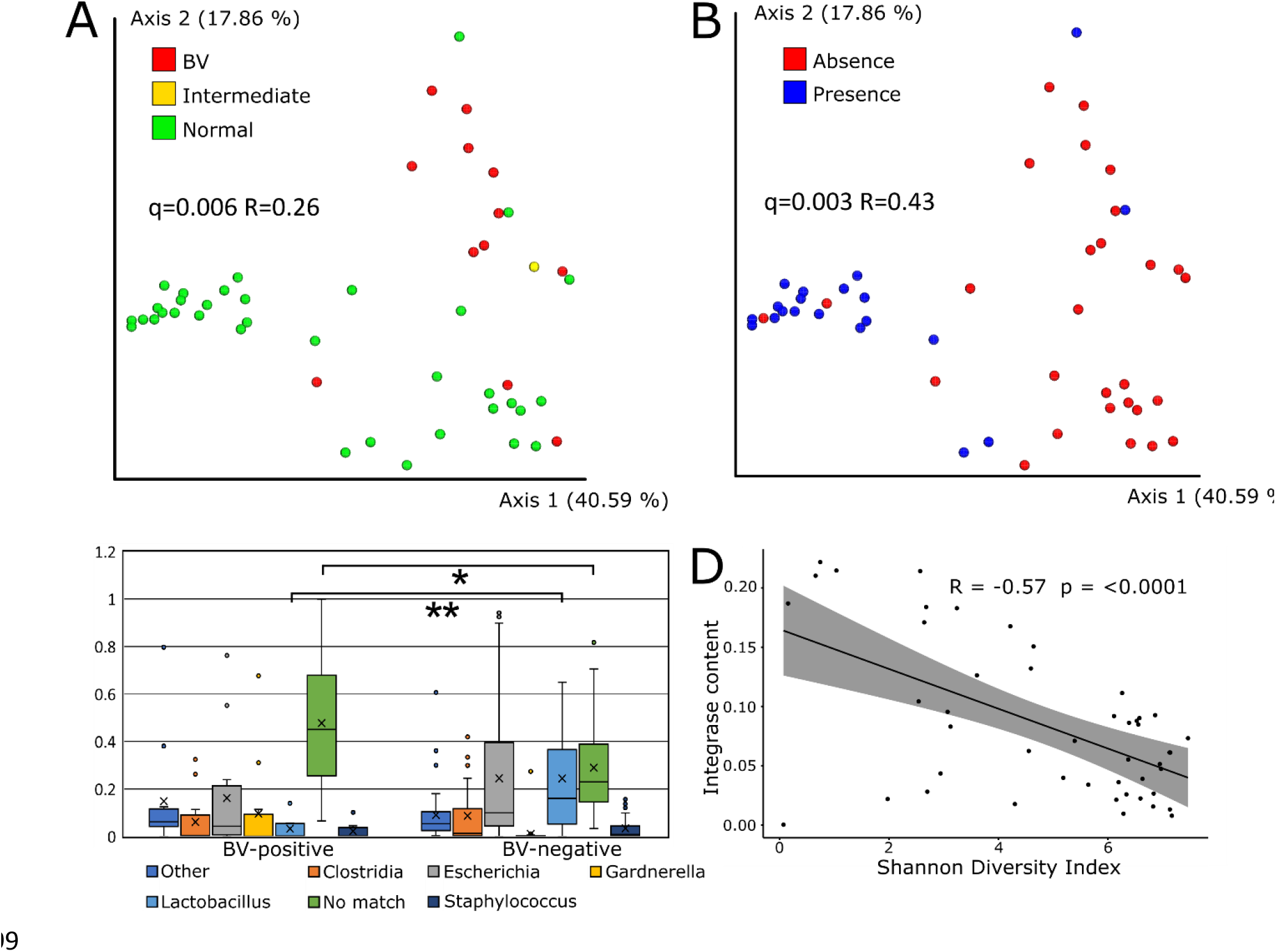
(A) Vaginal virome composition (Bray Curtis dissimilarity metric) varies with BV-status and (B) Lactobacillus crispatus presence/absence (determined by qPCR). (C) Relative abundance of WISH host genus predictions of vOTUs by BV status. Significance was calculated using Kruskall-Wallis Test (*=p<0.05, **= p<10^−3^). (D) Scatterplot plot showing Shannon diversity against the percentage of reads mapping to integrase genes by sample. Significance was calculated using the Pearson correlation.

### Sequence-based host prediction and integrase content

Using WIsH[30] to predict the bacterial host of viral OTUs showed that *Clostridium* was the most common predicted host genus, followed by *Lactobacillus* and *Gardnerella* (**Figure S4A**). BV-negative samples contained a significantly higher relative abundance of vOTUs predicted to have *Lactobacillus* spp. as their most likely host (p < 0.0001), while BV-positive samples contained significantly more vOTUs with no high-confidence host match (p = 0.03) (Figure 2C).

Lysogenic phages can be identified using marker genes such as integrases which are specific to lysogenic phages [36]. The ratio between lytic and lysogenic phages was estimated by comparing the integrase gene content between samples. There was no statistically significant difference in the fraction of reads mapping to integrase/transposase sequences between samples with different BV-status, although BV-negative samples had a higher mean fraction (0.091±0.004) compared to BV-positive samples (0.066±0.005) **(S4B)**. Interestingly, there was a strong negative correlation between the viral alpha diversity (Shannon diversity index) and the fraction of viral reads matching integrase genes in each sample (p-value < 0.0001) (Figure 2D).

### Bacteriophage-bacteria interactions

Co-abundance analysis of bacterial OTUs and vOTUs was performed using Regularized extension of Canonical Correlation Analysis (rCCA) to reveal correlations between individual phages and bacteria. The rCCA analysis was separately performed for BV positive and negative samples under the assumption that the microbial dynamics and environmental conditions such as pH and would be highly different between the two groups. The correlation analysis revealed large sets of strongly correlated bacterial OTUs and vOTUs as shown for the BV-positive samples in Figure 3. When comparing cross validation (CV) scores, BV-positive samples had strong correlations between bacterial and viral OTUs (CV-score 0.82) in comparison to BV-negative (CV-score 0.38) (**Figure S5A**) and all 48 samples together (CV-score 0.36) (**Figure S5B**).

**Figure 3.**
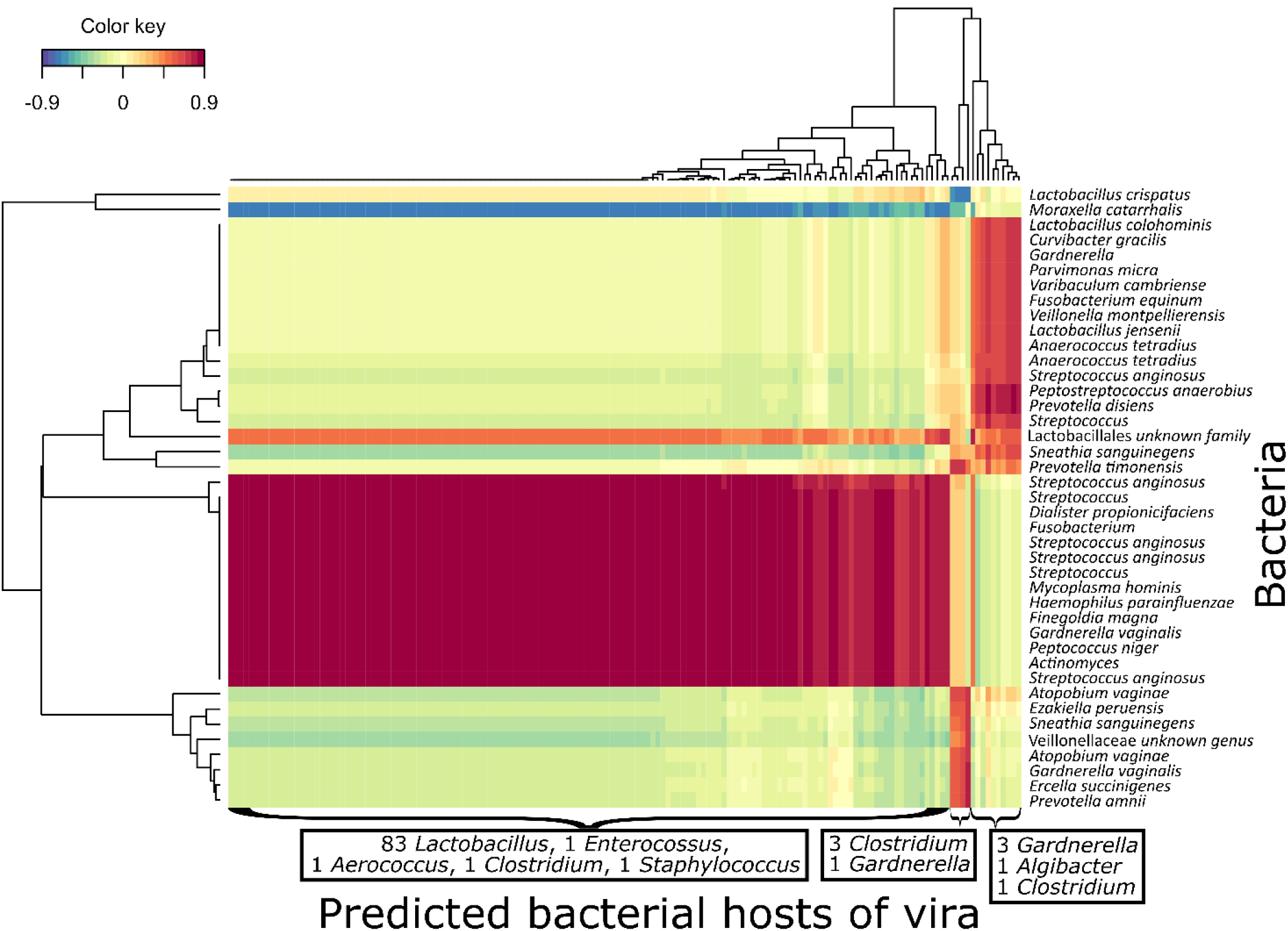
Clustered Image Map (CIM) of regularized Canonical Correlation Analysis (rCCA) between relative abundances of bacterial OTU and viral OTUs. Colour grade shows strength of correlation between individual bacterial and vOTUs. Viral clusters were summarized based on their bacterial host genus as predicted by WIsH as only a minority had matches in viral databases. Bacterial OTUs that have several entries had distinct 16S sequences and are possibly different strains. Correlations above 0.7 are shown. CV-score = 0.82.

## Discussion

### Main findings

This study provides the first in-depth characterization of the vaginal DNA virome based on virus like particle purification followed by metagenomic sequencing and de-novo assembly, constructing a total of 773 vOTUs (of these 302 represent previously undescribed viruses/phages). We found that the composition of both the prokaryotic and the eukaryotic viral communities varied strongly between BV-negative and BV-positive samples. Further, clear co-abundance patterns between certain bacteria and vOTUs indicate that these two components of the vaginal microbiome are strongly interlinked. Interestingly, the eukaryotic viral component differed significantly between BV-positive and negative samples, even though these viruses are not directly interacting with the bacterial community.

The identified vOTUs were predominantly dsDNA viruses, unidentified viruses and a small proportion of ssDNA viruses, but only a minority had exact matches in viral databases. This is not unexpected considering the lack of previous characterization of the vaginal virome using de-novo assembly and an overall underrepresentation of viral genomes in relevant databases[37].

Co-abundance correlation showed that groups of mainly anaerobic bacteria including *G. vaginalis* and *Moraxella catarrhalis* are negatively correlated with a large number of vOTUs predicted to target *Lactobacillus* spp. and other commensal bacteria. This is likely to reflect that a vaginal microbiota that contains many lactobacilli have low abundances of BV-associated bacteria, and therefore also lack the corresponding phages. In this aspect the finding of distinct viral groups could indicate the presence of distinct vaginal viral communities, or viral CSTs, that correspond to distinct bacterial CSTs [38]. Moreover, viral community structure was strongly correlated with presence (determined by qPCR thresholds) of key beneficial bacterial species (*L. crispatus, L. iners*) as well as with known pathogens (*G. vaginalis*, *A. vaginae*). This demonstrates a clear link between the viral component and key bacterial indicators of vaginal microbiome health and dysbiosis.

The observation that the eukaryotic viral community differed by BV-status was surprising as these are not directly linked to the bacterial community. It can be speculated that rather than being directly linked to the dysbiotic bacterial community associated with BV, the differences in the eukaryotic viral community composition reflects environmental factors, such as low pH, which is linked with healthy microbiota status and has been shown to inhibit viral infectivity and survival[39]. In line with this, a *Lactobacillus*-dominated cervicovaginal microbiota has been associated with reduced genital HIV viral load and HIV/STI prevalence[40]. Of the identified eukaryotic viruses DNA, the Herpesvirales and Pappilomaviradae orders were the only groups currently known to be associated with human disease.

### Biological role of virome/phages in vaginal health and disease

Given the observation that large groups of bacteria and bacteriophages correlated in a biologically meaningful fashion it is indeed likely that bacteriophages play a role in shaping the vaginal bacterial community. Expansions of the viral community appear to be caused by externally originating lytic phages rather than activation of prophage elements from bacterial genomes, as samples with highly diverse viral communities contain a smaller ratio of lysogenic phages. The reason for this could be that a more diverse and unstable bacterial composition, favours the lytic lifestyle in the corresponding virome and that a simple, stable community favours temperate phages. This is supported by computational models showing how the virulent strategy works best for phages with a large diversity of hosts, and have access to multiple independent environments reachable by diffusion[41]. Therefore the vaginal mucosa, which is highly perturbed by multiple factors such as sex, scanning and menstruation[42], could favour lytic replication in comparison to the more sequestered intestinal mucosa where lysogeny dominates[43].

There are several possible mechanisms of phage-related vaginal dysbiosis. One is introduction of foreign phages (i.e. from the partner) depleting the commensal bacteria and allowing pathogen colonization. Alternatively, pathogenic bacteria could be directly colonizing with their corresponding phages as passengers. In the first case vaginal stability would require commensal bacteria to be resistant to invading phages, for example by containing a prophage element of a related phage, providing immunity in the form of superinfection exclusion[44]. In the scenario of direct pathogen colonization, phages in the vaginal mucosa could protect against these, thereby providing non-host derived immunity. Women lacking these protective phages in the vaginal mucosa would then be more susceptible to develop dysbiosis. In this case, phage-therapy with a cocktail of phages could be used to target vaginal pathogens specifically, allowing the commensal bacterial population to re-establish. From this study, it is not possible to determine whether bacterial dysbiosis preceded or followed changes in the viral community. Studies using longitudinal vaginal samples are needed to elucidate the temporal dynamics of these two communities in detail.

### Comparisons to other studies

Our findings contrast with previous findings of Gossman et al.[19], where no distinct viral community structure differences between BV-positive and negative samples were found. However, contrary to Gossman et al., the present study up-concentrated VLPs prior to sequencing and used a de novo construction approach allowing detection of viral sequences not already in the databases, significantly improving sensitivity.

### Limitations

Viral metagenome sequencing measures relative and not absolute abundance, it is therefore possible that there could be differences in overall viral load that were not detected. The relatively small samples size and lack of longitudinal samples limits advanced analysis of phage-bacteria dynamics. The limited detection of negatively correlated phage-host pairs is likely due to the lack of longitudinal sampling, as murine enteric studies have shown that the increase in phage abundance and host decrease occurs over a 7-8 day period before reaching new stable levels [45]. This study reveals only a smaller fraction of the eukaryotic viral component as the majority of eukaryotic viruses have RNA genomes[46].

### Conclusions

In conclusion, this first in-depth investigation of the vaginal DNA virome finds that the vaginal viral community is strongly correlated with the vaginal bacterial community and BV. Therefore, including the viral component has the potential to provide a much more complete understanding of the mechanisms in vaginal microbial health and dysbiosis. What remains to be determined is the strength and of equally importance, the direction of the interaction between the vaginal viral and bacterial component which will require longitudinal studies.

## Notes

### Funding

The study was partially funded by Christian Hansen A/S with a grant to DSN.

### Conflicts of interest

TH has received honoraria for lectures from Ferring and Merck. PH received unrestricted research grants from MSD, Merck, and Ferring as well as honoraria for lectures from MSD, Merck, Gedeon-Richter, Theramex, and IBSA. JSJ has received speaker’s fee from Hologic, BD and Cepheid and serves scientific advisory boards of Roche Molecular Systems, Abbott Molecular, and Cepheid. PH, TH and JSJ received a research grant from Osel inc. which produces LACTIN-V, a live biotherapeutic product with *Lb. crispatus*. PH and TH are listed as inventors in an international patent application (PCT/UK2018/040882) involving “Use of vaginal lactobacilli for improving the success rate of in vitro fertilization”.

### Disclaimer

The funders had no role in study design, data collection and interpretation, or the decision to submit the work for publication.

