## Supplementary figures and materials for "Characterization of the vaginal DNA virome in health and dysbiosis: an opening study in patients with non-female factor infertility"

### Supplementary materials

#### S1 Vaginal viruses: taxonomy, alpha and beta-diversity

##### A Relative viral abundance at group level

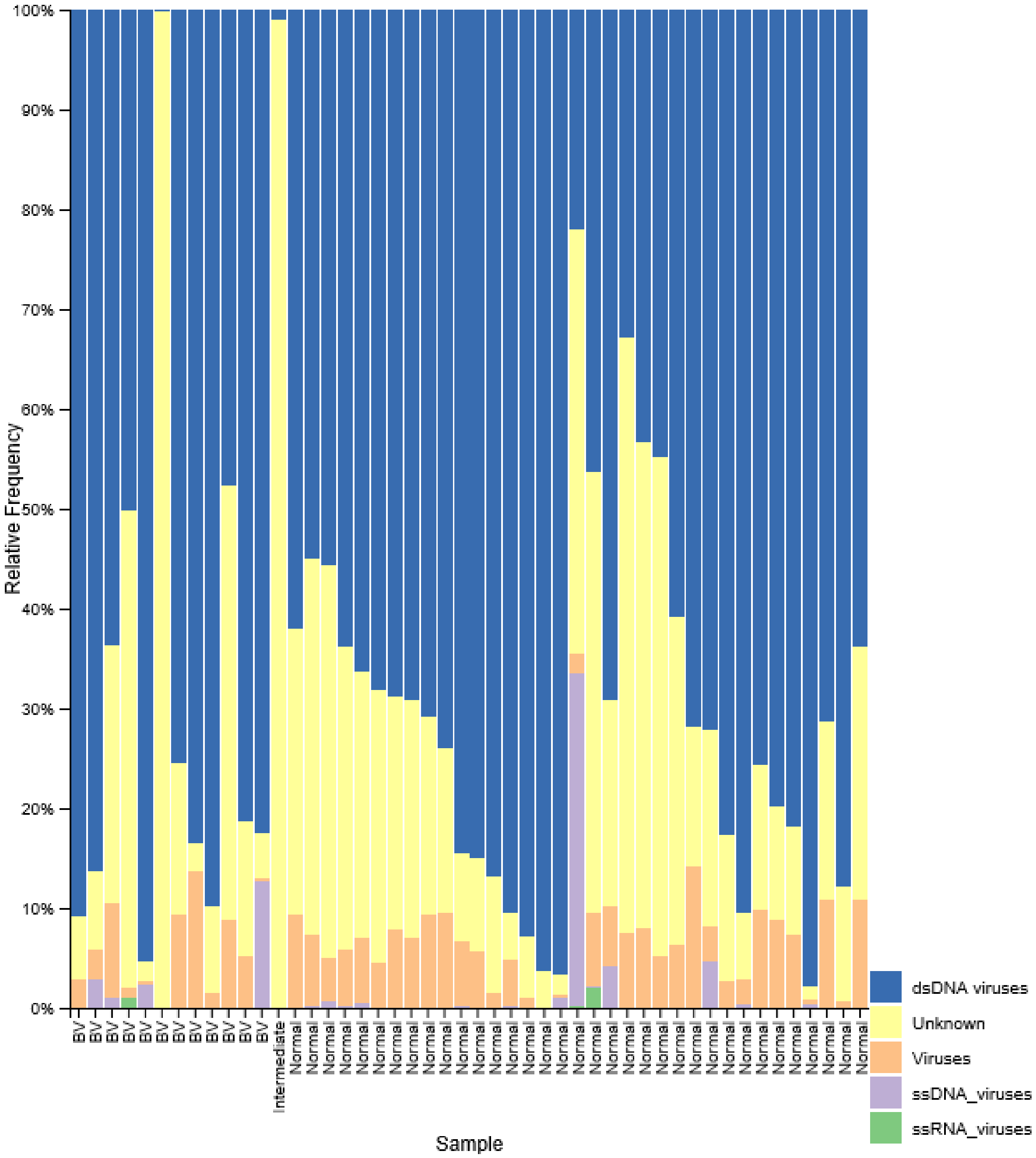


**Figure S1A** Relative viral abundance summarized at group level. Taxonomy based on closes viral database hit Samples grouped by BV-status as determined by nugent scoring.

#### B Viral alpha diversity – Shannon index

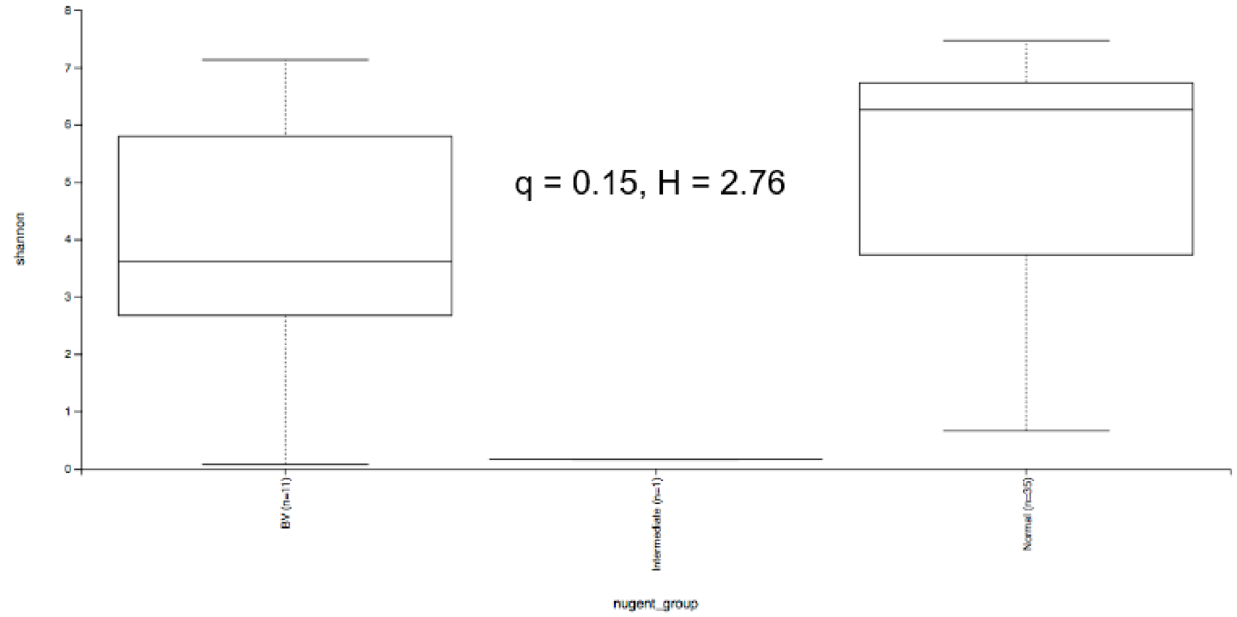


#### C Viral alpha diversity – Observed OTUs

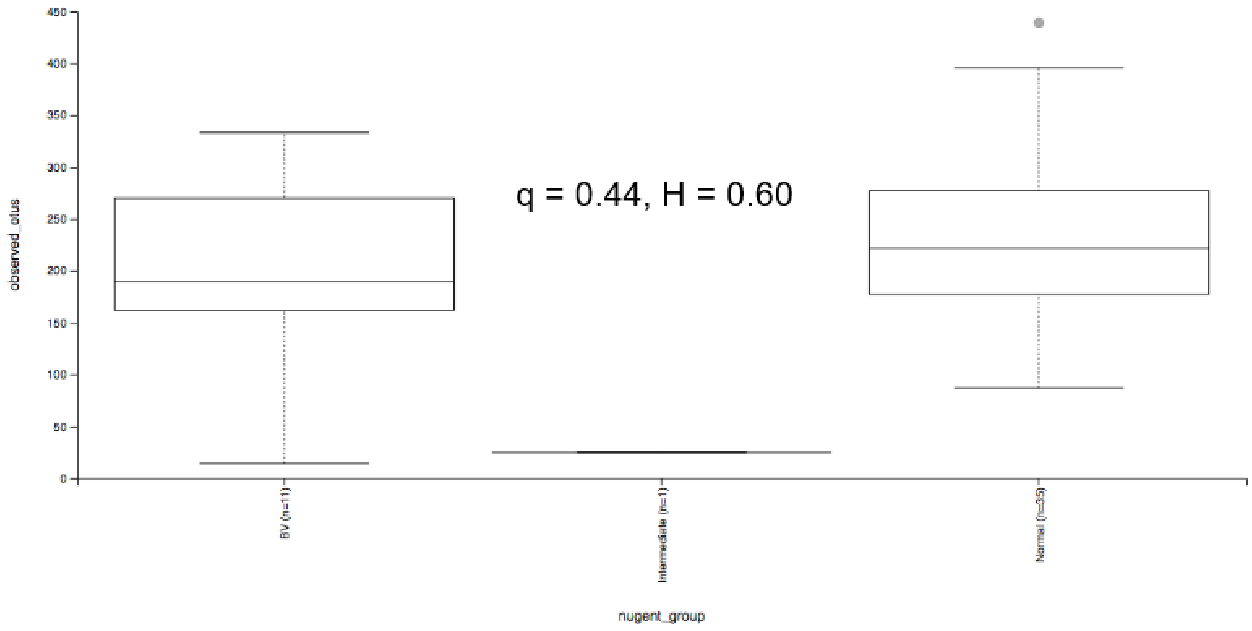


**Figure S2B,C.** Viral alpha diversity by (B) Shannon diversity index and (C) Observed OTUs. Significant correlations are marked with (*) tested by Pairwise Kruskall-Wallis test.

D Bacterial vs viral alpha diversity

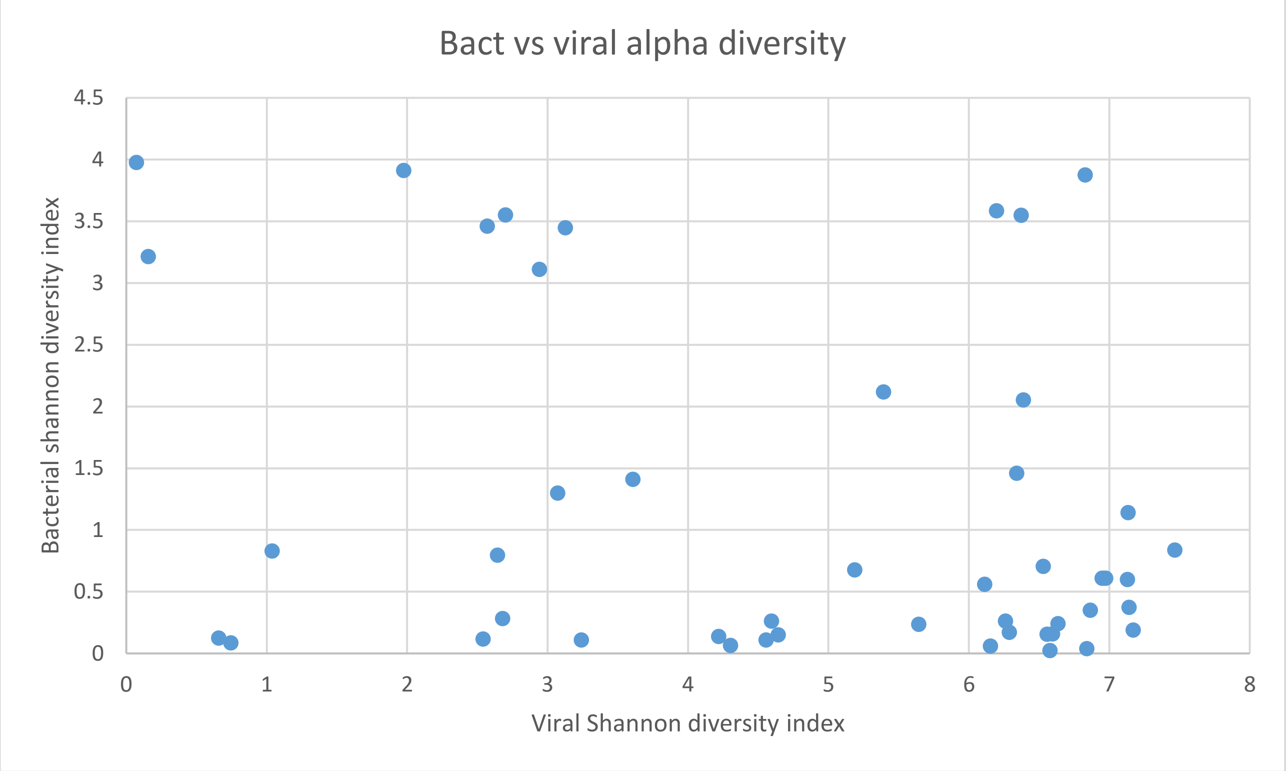


**Figure 2D.** Boxplot showing viral vs bacterial alpha diversity by Shannon diversity index. No significant correlation.

#### S2 Vaginal Eukaryotic viruses

##### A Relative eukaryotic viral abundance at highest possible level of taxonomic identification

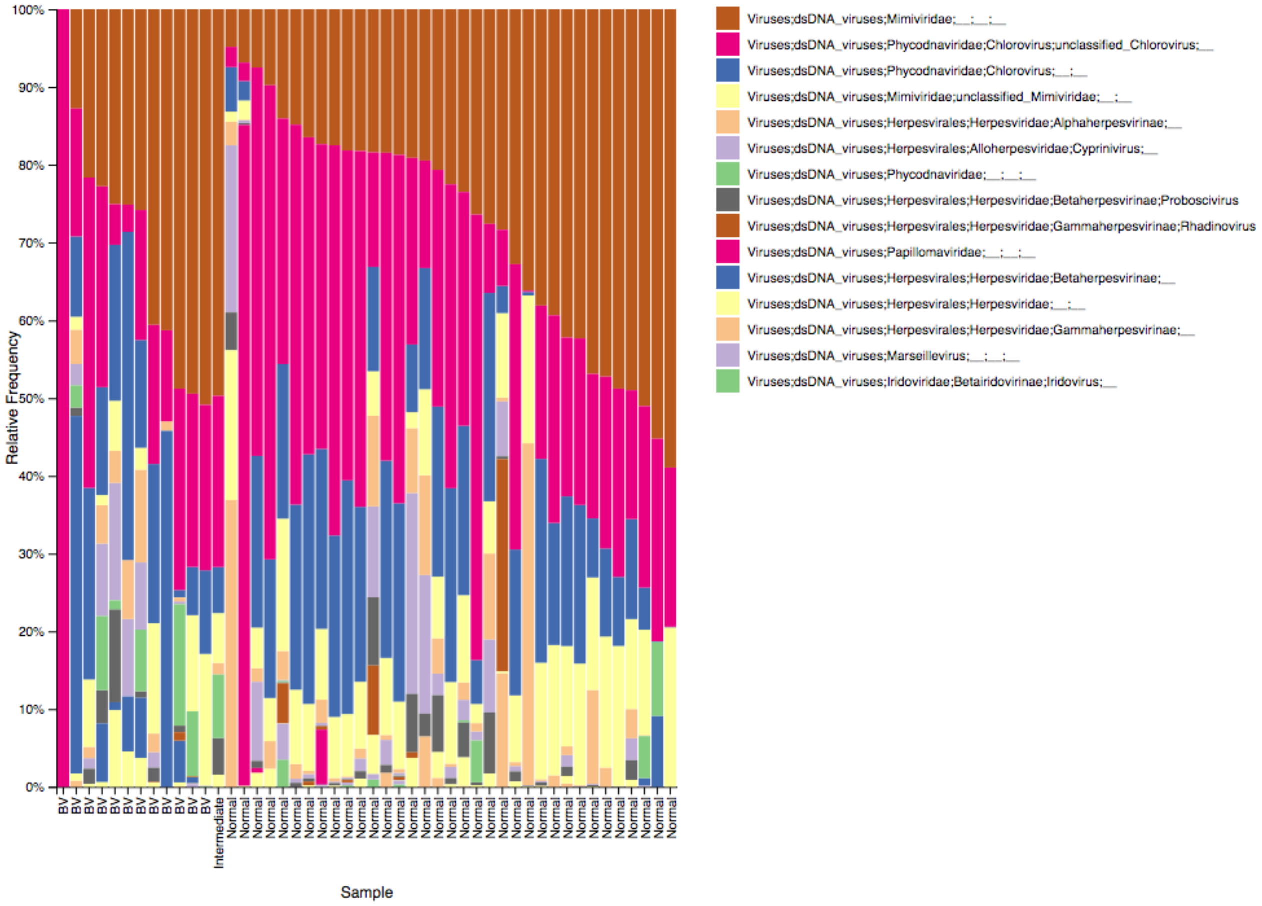


**Figure S2A** Relative abundance of eukaryotic viral OTUs. Only vOTUs with eukaryotic viral database matches were included. Taxonomic identification to highest possible level.

#### B Eukaryotic viral composition by BV-status Bray-Curtis dissimilarity index

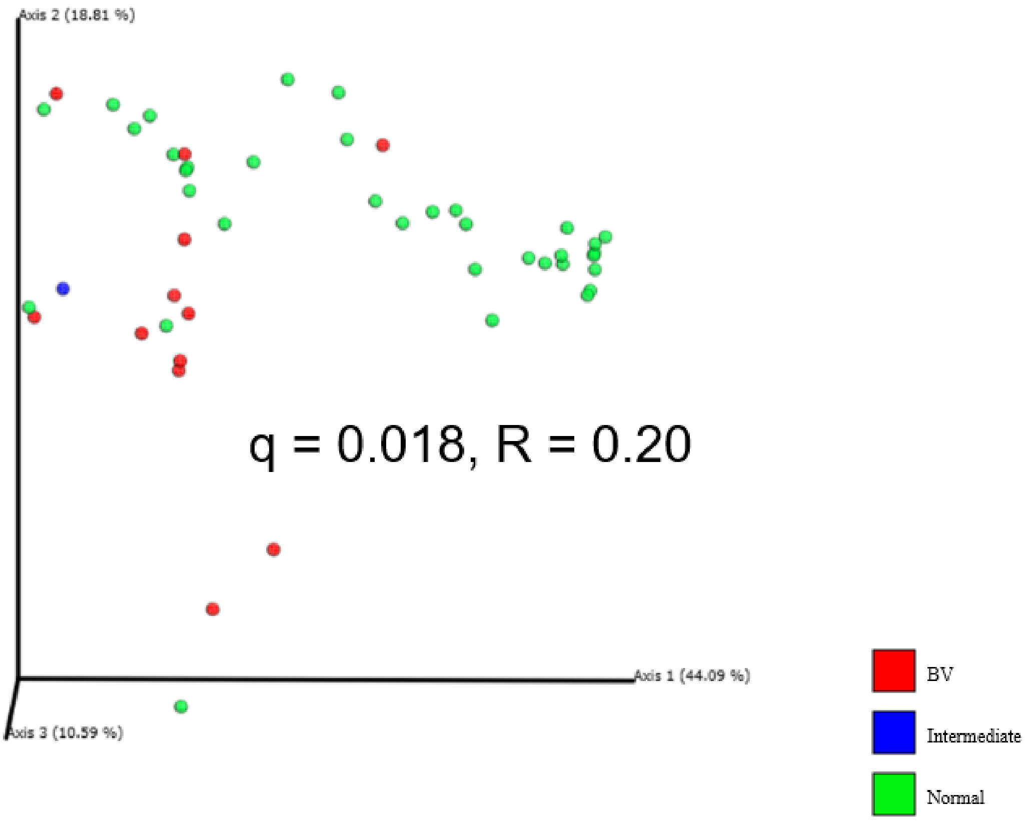


**Figure S2B** Bray-Curtis dissimilarity beta diversity by BV-status: PCoA and group comparison by pairwise ANOSIM

| Bacteria | R | p-value |
|---|---|---|
| *Lactobacillus crispatus* | 0.43069 | 0.001 |
| *Lactobacillus iners* | 0.148474 | 0.002 |
| *Lactobacillus jensenii* | -0.10973 | 0.987 |
| *Lactobacillus gasseri* | -0.114594 | 0.962 |
| *Gardnerella vaginalis* | 0.209314 | 0.003 |
| *Atopobium vaginae* | 0.247796 | 0.002 |

**Figure S2D** Bray-Curtis dissimilarity by presence/absence of key bacterial species as determined by qPCR. ANOSIM pairwise comparisons.

#### S3 Vaginal bacterial compoment

##### A Bray-Curtis dissimilarity beta diversity by BV-status: PCoA and pairwise anosim

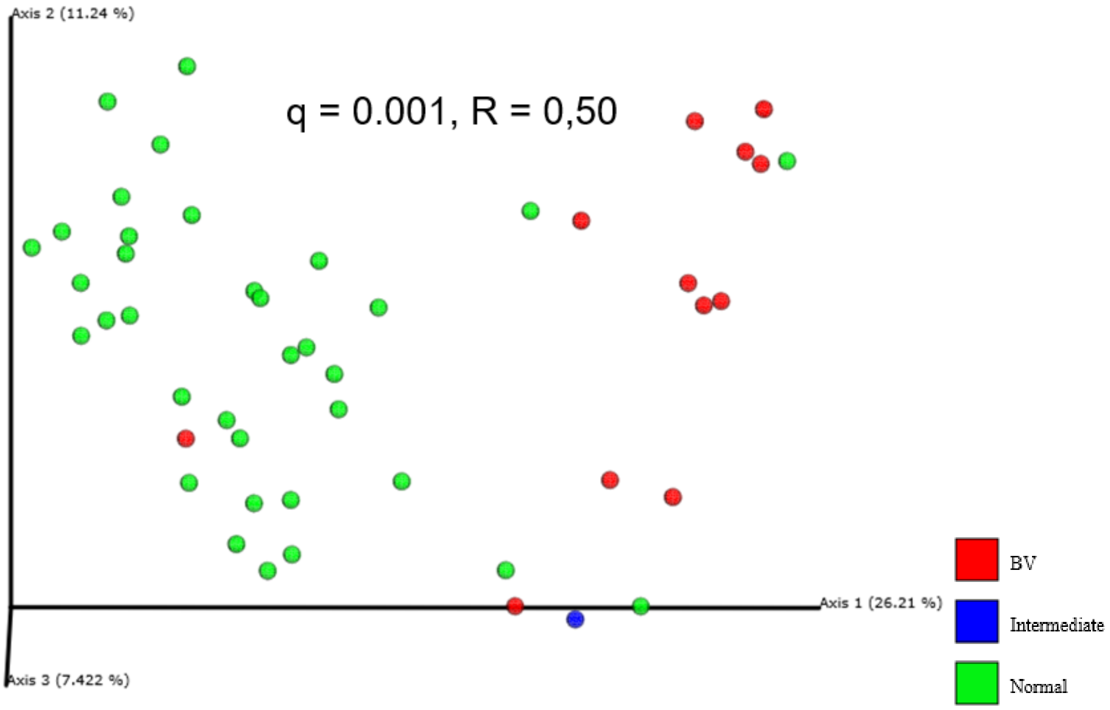


**Figure S2B** Bray-Curtis dissimilarity beta diversity of vaginal bacterial component(16S) by BV-status: PCoA and group comparison by pairwise ANOSIM

#### S4 Viral-bacterial correlations

##### S4A WIsH host prediction by relative abundance – all samples

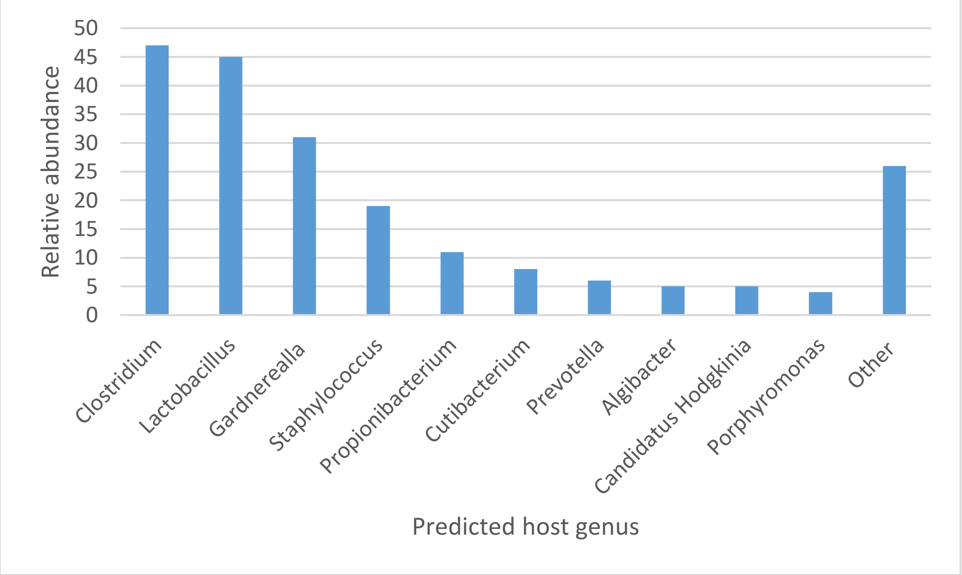

**Figure S4A** WIsH host prediction of vOTUs across all samples by relative abundance, grouped at genus level.

#### S4B Integrase content by BV-status

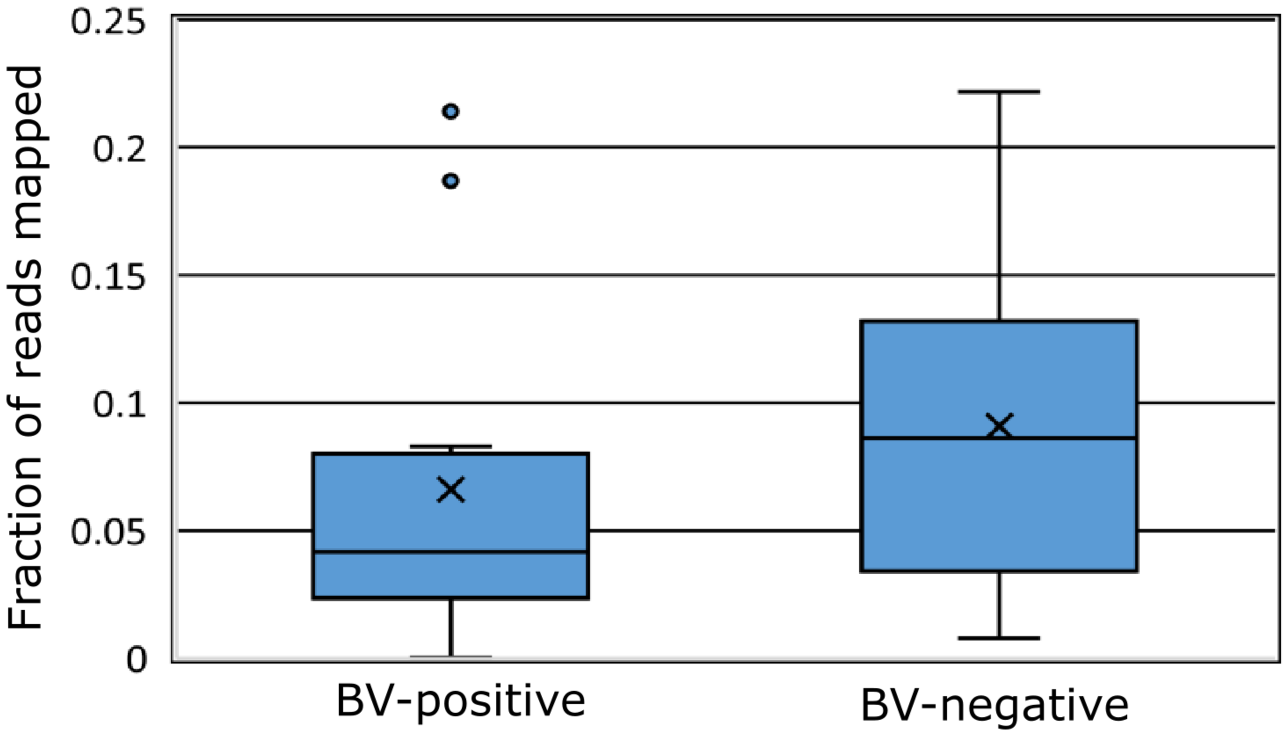


**S4B** Integrase content of viral component by BV status. Percentage of reads mapping integrase genes. No significant difference.

#### S5 Viral-bacterial correlation: Regularized extension of the Canonical Correlation Analysis

##### A All samples – Correlations above 0.4 shown

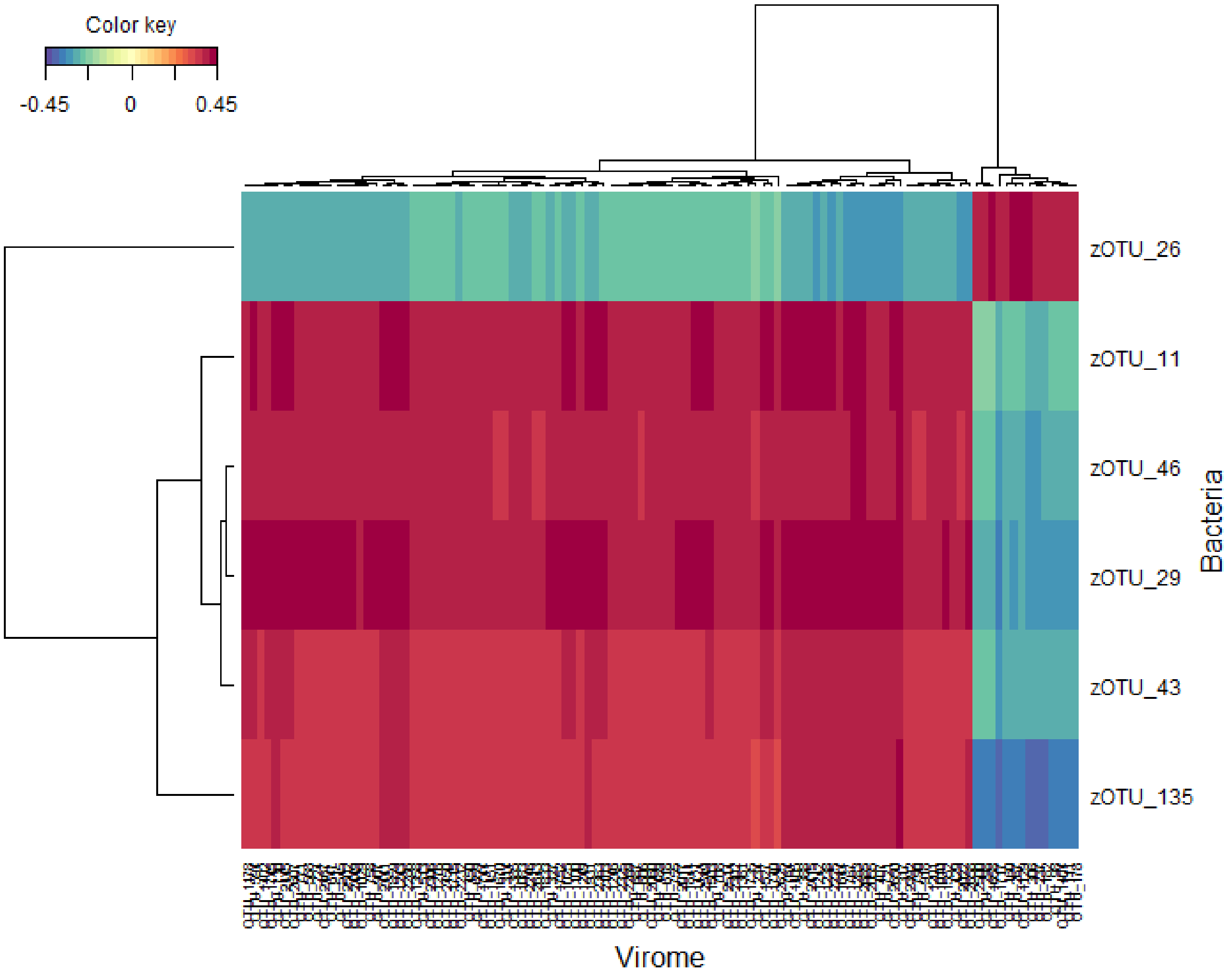


##### B BV negative samples – Correlations above 0.4 shown

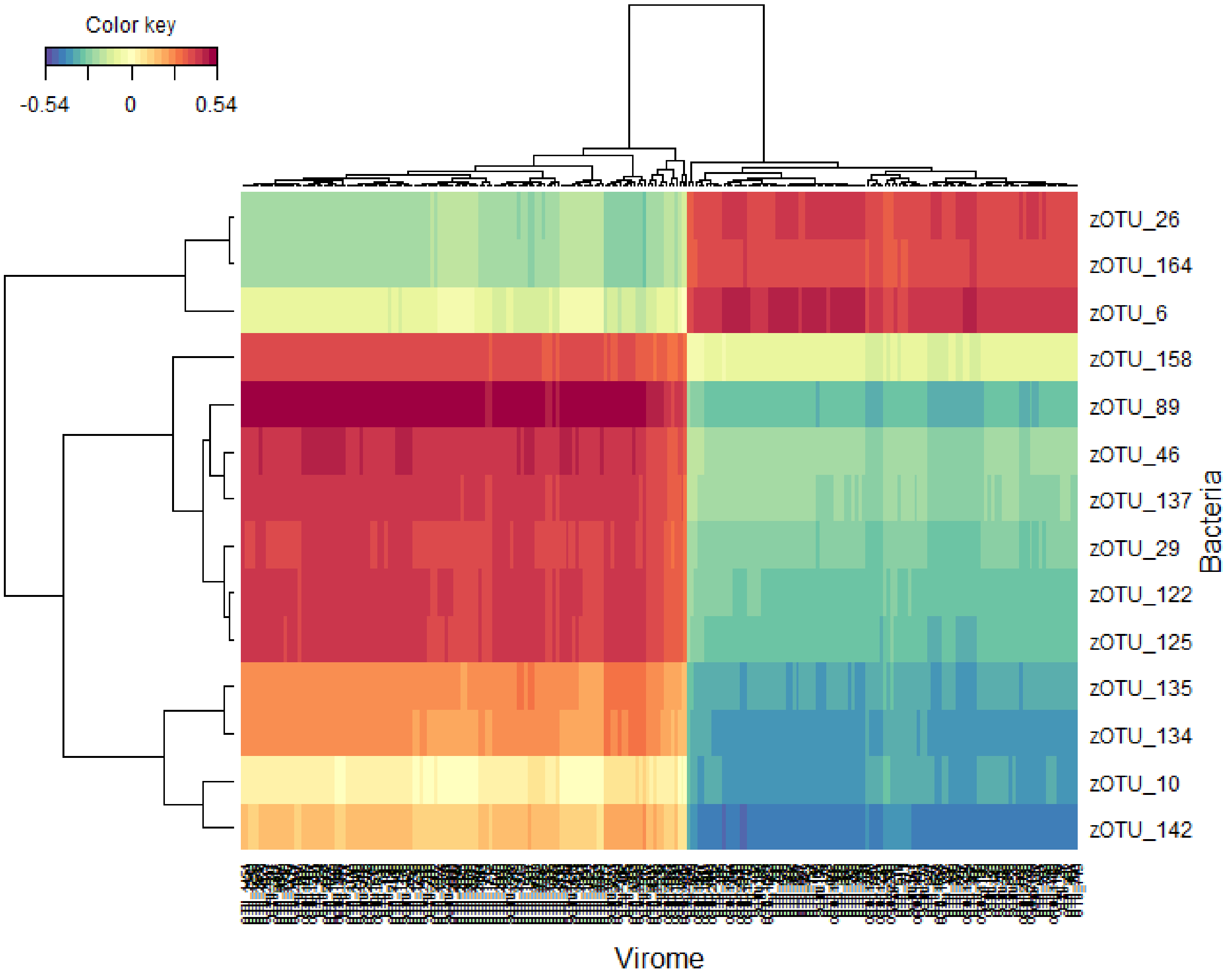


#### S6 Trimming, cleaning, assembly and deduplication of VLP-derived metagenome reads

The raw reads were trimmed from adaptors and barcodes using Trimmomatic v0.35[1] (>97% quality [seedMismatches: 2, palindromeClipThreshold: 30, simpleClipThreshold:10; LEADING: 15; MINLEN: 50], removed from ΦX174-control DNA and de-replicated (Usearch v10)[2]. Non-redundant high-quality reads with a minimum size of 50nt were retained. The presence of non-viral DNA was quantified using 50,000 random forward-reads from each sample, which were queried against the human genome, as well as all the bacterial and viral genomes hosted at NCBI using Kraken2 [3]. Similarly, reads were blasted against the non-redundant protein database available at UniProtKB/Swiss-Prot (-evalue 1e-3, -query_cov 0.6, -id 0.7), the ribosomal 16S rRNA (GreenGenes 13_5[4]) and 18S rRNA (Silva, release 126[5]) databases (-evalue 1e-3, -query_cov 0.97, -id 0.97).

Reads generated from VLP-derived DNA sequencing were subjected to within-sample de novo assembly. Assembly was carried out using metaSPAdes v3.5.0[6] and only scaffolds (here termed "contigs") with a minimum length of 1,000 nt were retained. Contigs generated from all samples were pooled and de-replicated by multiple blasting and removing those contained in over 90% of the length of another (90% similarity), as outlined previously[7]. To check the presence of non-viral DNA contigs, de-replicated contigs were evaluated according to their match to a wide range of viral proteins, [viral non-redundant RefSeq [8], virus orthologous proteins (www.vogdb.org), and the prophage/virus database hosted at PHASTER (www.phaster.ca[9])], reference independent k-mer signatures [VirFinder[10]], viral genomes RefSeq [NCBI, Kraken2] as well as their match to bacterial (--confidence 0.08), plant (Kraken2, --confidence 0.3) and human genomes (Kraken2, --confidence 0.1) deposited in the NCBI database. All contigs that did not match any database or that matched viral sequences, viral proteins or viral k-mers were subsequently retained and categorized as viral contigs.
